# Comparison of lumbar paraspinal muscle size and density between members of the US Framingham Heart Study and Tsimane forager-horticulturalists of Bolivia

**DOI:** 10.64898/2026.09.25.754176

**Authors:** Benjamin Sibson, Brett Allaire, Brigitte Holt, Fjola Johannesdottir, Shreyas Durga, Daniel Cummings, Michael Gurven, Hillard Kaplan, Benjamin Trumble, Jonathan Stieglitz, Mary Bouxsein

**Affiliations:** Endocrine Unit, Massachusetts General Hospital, Boston, MA, USA; Department of Orthopaedic Surgery, Harvard Medical School, Boston, MA, USA; Center for Advanced Orthopaedic Studies, Beth Israel Deaconess Medical Center, Boston, MA, USA; Department of Anthropology, University of Massachusetts Amherst, Amherst, MA, USA; Department of Anthropology, University of New Mexico, Albuquerque, NM, USA; Economic Science Institute, Argyros School of Business and Economics, Chapman University, Orange, CA, USA; Department of Anthropology, University of California, Santa Barbara, CA, USA; Center for Evolution & Medicine, School of Human Evolution and Social Change, Institute of Human Origins, Arizona State University, Tempe, AZ, USA; Toulouse School of Economics, Université Toulouse 1 Capitole, Toulouse, FRA

## Abstract

Paraspinal muscles support and move the spine, and reduced size and density of these muscles is associated with increased risk of vertebral fracture and back pain. We test the hypothesis that subsistence-oriented forager-horticulturalists (Bolivian Tsimane) have larger and denser paraspinal muscles, and smaller age-related differences in paraspinal muscle size and density, than more sedentary post-industrialized suburbanites (Framingham Heart Study [FHS]). We measured cross-sectional area (CSA) and density of trunk extensors and psoas major at L3 from computed tomography scans of 411 FHS and 330 Tsimane (50% women, aged 50–90 years). Sex-stratified linear regressions determined associations of muscle CSA and density with age and cohort (Tsimane or FHS [ref.]), with and without an age–cohort interaction term. Covariates were height and weight. After adjusting for covariates, Tsimane women had 11–14% larger and denser extensors, and 17–26% smaller and less-dense psoas major, than FHS women. Tsimane men had 8–21% smaller and less-dense psoas major than FHS men. The age-related difference in extensors density was 41% smaller for Tsimane men than for FHS men. There were no other significant age-related cohort differences in muscle CSA or density. We observed a different balance in the size and density of trunk extensors versus psoas major between Tsimane and FHS. A possible explanation for this observation is cohort differences in the amount and types of physical activity, although future work is required to determine the influence of lifestyle and environmental factors on paraspinal muscle size and density.

## Introduction

Paraspinal muscles run vertically adjacent to the spine and support and move the spine during postural and locomotor behaviors [1,2]. Reduced paraspinal muscle size can impair physical function (e.g., balance) and increase risk of back pain and vertebral fractures [3–6]. Reduced paraspinal muscle density, usually reflecting higher fat infiltration within the muscle, is also associated with impaired balance and vertebral fractures, independent of muscle size [7,8]. Together, paraspinal muscle size and density (here referred to as “muscle morphology”) determine the amount of contractile tissue within a muscle and therefore its capacity to generate force and do work. Posterior paraspinal muscles extend the trunk (i.e., “extensors”), and include erector spinae, transversospinalis, and quadratus lumborum. The anterior paraspinal muscle— psoas major—flexes the trunk and hips [9]. It is useful to distinguish between extensors and psoas major because their morphologies are independently associated with vertebral fracture risk [10–14].

Musculoskeletal conditions of the spine, such as back pain and vertebral fractures, become more prevalent with advancing age; they are also the most common type of musculoskeletal condition [15–18]. The high prevalence of spine conditions is due to several factors. The recent trend of industrialization changed many people’s exposomes (i.e., lifestyle and environmental exposures) in ways that may negatively impact spine health [19]. For example, increased sedentariness may cause people to develop weak, fatigable paraspinal muscles that poorly support the spine, potentially resulting in injury [20–22]. More generally, it has been argued that the evolution of habitual bipedal posture may increase human susceptibility to spine conditions. Habitual bipedal posture required humans evolve, among many other anatomical changes, an anterior curvature in the lumbar spine (“lordosis”); lordosis curvature creates shear forces in the lumbar spine that may increase risk of disc degeneration, spondylolisthesis, and other injury [23–26]. This combination of recent exposome changes and humans’ evolved spine anatomy likely contributes to the high prevalence of spine conditions.

Paraspinal muscle size and density typically decline alongside the increasing prevalence of spine conditions with aging; prior work shows decreases of 8–17% per decade among US adults over age 40 years [27]. These age-related declines may be caused by several factors, the most salient being cellular senescence and reduced physical activity (and associated mechanical loading) [28,29]. The percentage of US adults meeting national physical activity recommendations decreases by almost a third from ages 18 to 65 years [30]. Reduced physical activity can affect muscle morphology because the mechanical loads induced by physical activity create microtears in muscle fibers that lead to subsequent hypertrophy and growth [31]. In the absence of such mechanical loads, muscle atrophies.

The phenomenon of muscle atrophy may be due to evolutionary selection for energy conservation. Muscle tissue is metabolically costly to develop and maintain [32,33]. According to energy allocation theory, muscle atrophy would be beneficial because having less muscle tissue to maintain makes more energy available for other vital functions, such as storage, growth, reproduction, and maintenance of other tissues [34–36]. A related consideration is the argument as to whether human energy expenditure is constrained. In an energy-limited context, higher energy expenditure on physical activity may actually compromise metabolic investment to muscle at the expense of other essential bodily functions; the result would be less muscle repair and growth [37]. An alternative hypothesis is that physical activity-induced muscle damage will still preferentially allocate energy towards muscle repair and hypertrophy—and slow cellular senescence—possibly at the expense of other bodily systems and functions [38].

To move towards a better understanding of age-related change in paraspinal muscle morphology, it is useful to compare populations with different exposomes. Most studies of muscle morphology, however, involve only post-industrialized populations [39]. Non-industrialized subsistence-oriented populations typically experience more variable and extreme exposomes than post-industrialized populations [40–42]. For example, a study of over 500 Bolivian Tsimane adults—forager-horticulturalists who subsist by farming, hunting, gathering, and fishing in the Amazon basin—showed that both women and men engage in 175 min/d of moderate-to-vigorous physical activity (MVPA) and walk 16,000 steps/d [43]. In comparison, a study of over 2,000 adults of the US-based Framingham Heart Study showed they engage in 20 min/d of MVPA and walk less than 8,000 steps/d. Post– and non-industrialized populations differ not only in the amounts, but also in the types, of physical activity they habitually perform. For example, starting early in life, Tsimane hunt, collect water and firewood, garden, fish, and carry children with the hands [44,45]. Most of these behaviors involve high levels of back muscle activity and vertebral loading [22]. Given different exposomes between post– and non-industrialized populations, especially physical activity, it is reasonable to expect population-level differences in paraspinal muscle morphology.

The aim of this observational study was to compare average and age-related differences in paraspinal muscle size and density between post-industrialized and non-industrialized population-based cohorts. The post-industrialized cohort were members of the US Framingham Heart Study (FHS) [46]. The non-industrialized cohort were Tsimane participants in the Tsimane Health and Life History Project (THLHP) [47]. We tested the hypothesis that the average size and density of Tsimane trunk extensors and psoas major are greater, and age-related differences are lower, than FHS. Higher physical activity-induced mechanical loading is associated with larger and denser paraspinal muscles [48,49], suggesting a Tsimane bias due to their high physical activity; FHS are more sedentary, predicting lower muscle size and density and greater age-related differences. Our alternative hypothesis was that the average size and density of Tsimane extensors and psoas major are lower, and age-related differences are greater, than FHS. This hypothesis followed from the notion that high energy expenditure among Tsimane (i.e., from high physical activity, high pathogen burden [50]) may compromise the growth and maintenance of their paraspinal muscles.

## Methods

### Study cohorts and participants

The THLHP has studied Tsimane since 2002 [47]. Teams of scientists, clinicians, and local research assistants visit Tsimane villages regularly and perform clinical examinations to study multiple aspects of Tsimane exposome and health. The cross-sectional sample of this study includes people who self-identified as Tsimane, were aged between 50–90 years, and underwent an abdominal computed tomography (CT) scan in 2024–25 (*N* = 330; 50% women) as part of a broader initiative to study chronic diseases of aging, including musculoskeletal conditions, cardiovascular disease, and dementia. The FHS has studied three generations of US adults since 1948 [51]. The cross-sectional sample of this study includes people from the Offspring and Third Generation cohorts that underwent abdominal CT scans in 2008–2011 (*N*=2,109) [46]. After categorizing the CT sample by sex and age decades 50–59, 60–69, 70– 79, and over 80 years, we selected a subsample participants to match the sex and age distribution of the Tsimane sample. *N*=411 FHS women and men aged 50–90 years (50% women) were included in this study. This study used previously collected, de-identified data. All study participants gave informed consent; for Tsimane, informed consent was also obtained from the Tsimane government that oversees research projects and from village leadership. All data collection protocols were approved by the IRB committees of Boston University, Beth Israel Deaconess Medical Center, the University of New Mexico Medical School, University of California Santa Barbara, and Universidad Mayor de San Simón Facultad de Medicina.

### Computed tomography scan acquisition

Tsimane underwent chest, abdominal, and pelvis CT scans at the Hospital Presidente German Busch in Trinidad, Bolivia. The CT scanner was a 16-detector row scanner (GE Brightspeed, GE Healthcare, Milwaukee, WI, US) with the following scan settings: a tube voltage of 100 kVp, tube current of 10–300 mA, 0.5 mm slice thickness, and gantry rotation of 500 ms. Tsimane CT scans were supervised and reviewed by at least one cardiologist and radiologist. Breath-hold instructions were given in the Tsimane language to minimize respiratory motion artifact and misregistration. FHS abdominal CT scans were acquired using a 64-slice PET/CT scanner (GE Discovery VCT, GE Healthcare, Milwaukee, WI, US) with the following scan settings: a tube voltage of 120 kVp, tube current of 300/350 mA (≤ 220/> 220 lb. body weight), 5 mm slice thickness, and gantry rotation of 350 ms [46]. Age was recorded and height and weight were measured at the time of the CT scans.

### Lumbar paraspinal muscle size and density

We measured paraspinal muscle size and density at the mid-plane of the third lumbar vertebrae (L3). L3 is the most common landmark used in CT imaging of paraspinal muscles, supports upper body weight, is the apex of lumbar lordosis, and is a target of osteophytes and other spine pathology [52–55]. We measured the trunk extensors—including erector spinae, transversospinalis, and quadratus lumborum—and psoas major. TotalSegmentator, a deep learning neural network framework validated for accurate and automatic segmentation of muscle tissue on CT scans, contoured extensors and psoas major at the mid-plane of L3 [56,57]. We measured muscle size as the cross-sectional area (CSA, mm^2^) within the muscle contours, averaging the right and left sides (Figure 1). We measured muscle density as the mean of voxel attenuation in Hounsfield units (HU) within the muscle contours, averaging the right and left sides with weighting according to muscle CSA. We excluded voxels outside the range of –50 to 150 HU before calculating CSA and density to remove voxels of pure fat, tendon, and bone along the periphery of the muscle contours.

**Figure 1.**
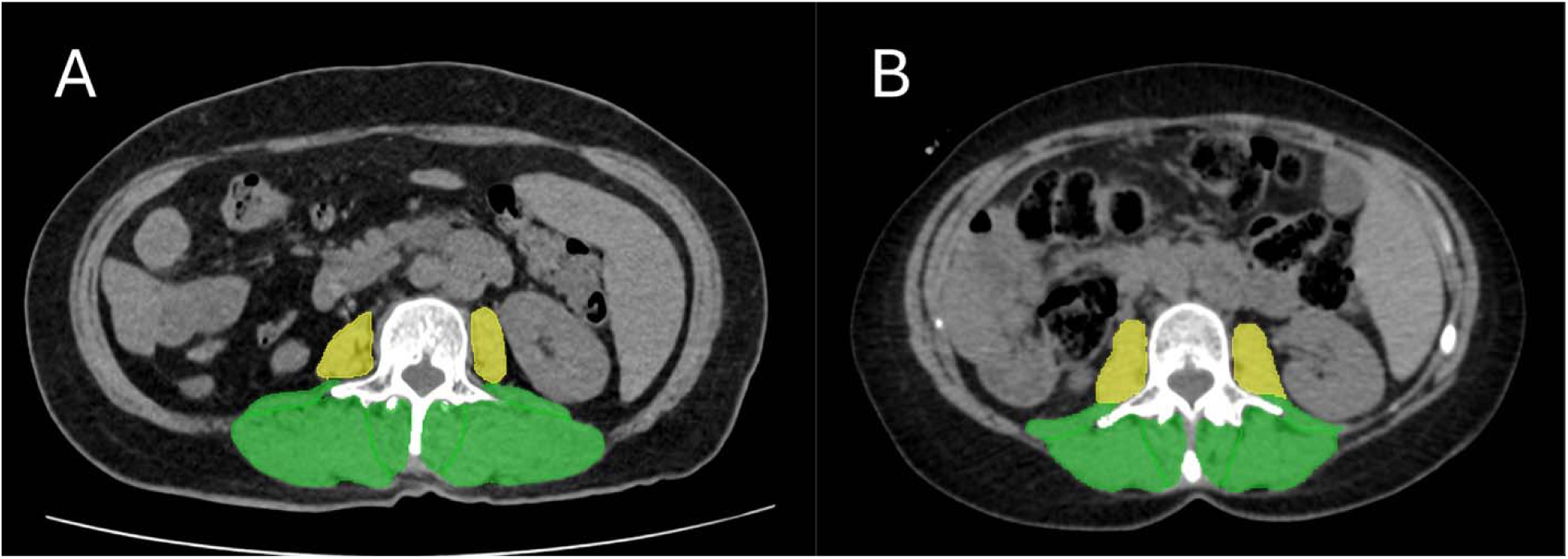
Computed tomography transverse images at the mid-plane of L3 for two age-matched (56 year-old) women across populations: (A) Tsimane woman who is 152 cm tall and weighs 61 kg, and (B) Framingham Heart Study woman who is 162 cm tall and weighs 68 kg. Extensors are shaded green, and psoas major is shaded yellow. Images in A and B have the same Field of View dimensions. Note the relatively larger extensors cross-sectional area (CSA) and smaller psoas major CSA for the Tsimane woman in (A).

To evaluate the accuracy of automated muscle segmentations in our sample, we compared CSA and density generated from manual muscle segmentations to CSA and density generated from automated segmentations in a subset of FHS (*N* = 272) and Tsimane (*N* = 40). Manual measurements of muscle have been used previously in large-scale studies to reliably examine paraspinal muscle morphology [7,27,58,59]. We linearly regressed manual versus automated measurements and created Bland-Altman plots of CSA and density for extensors and psoas major. Results support the validity of automated muscle measurements in this study: *R^2^* values and regression slopes were near unity, and each Bland-Altman plot showed a random distribution with no evidence of systemic bias (all 95% CI intervals included zero) (Figure S1–4).

### Statistical analysis

Sex-stratified cohort comparisons of age and anthropometrics (height, weight, and body mass index [BMI]) were made using Welch’s Two Sample *t* tests. Sex-stratified multivariate linear regression models quantified associations between muscle CSA or density with age and cohort (Tsimane vs. FHS [ref.]). In separate models, we also included an age–cohort interaction term to test for age-related differences in muscle CSA or density between cohorts. Covariates were height and weight. Height and weight are positively associated with paraspinal muscle CSA and skeletal muscle mass broadly [58,60]. Higher weight and BMI are also associated with more fat infiltration of paraspinal muscles, hence lower muscle density [61,62]. Therefore, to account for the potential effects of cohort differences in height and weight on paraspinal muscle CSA and density, we included height and weight as covariates in all regression models.

We checked linear regression assumptions of linearity, normality of residuals, heteroscedasticity, and influential outliers by inspecting residuals vs. fitted plots, q-q plots, scale-location plots, and residuals vs. leverage plots [63]. Models met assumptions of linearity and did not contain influential outliers. Heteroscedasticity was present in most regression models. We used Weighted Least Squares to minimize a weighted sum of the squared residuals in the regression models; this reduced heteroscedasticity to an appropriate level, based on re-examination of scale-location plots. Most q-q plots showed that residuals followed a normal distribution. Some q-q plots showed minor deviations at the extreme ends; because all regression models in this study had *N* ≥ 370, these minor deviations are robustly handled by the Central Limit Theorem. We also checked regression models for multicollinearity and found that all adjusted Variance Inflation Factors were < 2.

To determine mean differences in paraspinal muscle CSA or density between cohorts while accounting for the effects of age and body size, we used the linear regression models to calculate estimated marginal means of CSA or density, adjusting for the effects of age, height, and weight. To determine age-related differences in paraspinal muscle CSA or density between cohorts while accounting for the effects of body size, we used the linear regression models that included an age–cohort interaction term to calculate the estimated marginal effects of age on CSA or density for each cohort, adjusting for height and weight. Pairwise comparisons evaluated cohort differences in estimated marginal means and the estimated marginal effects of age on CSA or density. We applied Tukey’s HSD adjustment for multiple comparisons and performed analyses using R Studio Version 2023.09.1+494 (Posit Software, PBC, Boston, MA, USA). *P* values ≤ 0.05 were considered statistically significant, and we interpreted results based on effect size and precision, alongside statistical significance.

## Results

### Tsimane were shorter and lighter than FHS

Tsimane were 8–9% shorter, weighed 29–37% less, and had 13–18% lower BMI than FHS (Table 1, all *p*<0.001). Descriptive statistics for paraspinal muscle CSA and density values unadjusted for covariates, by sex and cohort, are found in Table S1.

**Table 1.** Descriptive statistics of age and anthropometrics for Tsimane and Framingham Heart Study (FHS).

| Variable | Women (N = 370) |  |  | Men (N = 371) |  |  |
| --- | --- | --- | --- | --- | --- | --- |
|  | Tsimane<br>N = 165 | FHS<br>N = 205 | Tsimane vs. FHS<br>% Diff | Tsimane<br>N = 165 | FHS<br>N = 206 | Tsimane vs. FHS<br>% Diff |
| Age (yrs) | 62 ± 8 | 63 ± 10 | — | 63 ± 8 | 64 ± 10 | — |
| Height (cm) | 149.1 ± 5.2 | 162.0 ± 5.7 | **−8.3 \*\*\*** | 159.9 ± 5.1 | 175.7 ± 6.3 | **−9.4 \*\*\*** |
| Weight (kg) | 54.9 ± 10.8 | 73.7 ± 16.5 | **−29.2 \*\*\*** | 62.0 ± 8.9 | 89.9 ± 14.9 | **−36.7 \*\*\*** |
| BMI (kg m<sup>−2</sup>) | 24.7 ± 4.4 | 28.1 ± 6.3 | **−13.0 \*\*\*** | 24.2 ± 3.1 | 29.1 ± 4.4 | **−18.2 \*\*\*** |

Data are Mean ± SD
\*\*\* $p < 0.001$ for Welch Two Sample $t$ test
% Diff calculated as the difference between Tsimane and FHS means, divided by their average, and multiplied by 100

### Tsimane women had larger and denser extensors than FHS women

Being younger, Tsimane, taller, and heavier were associated with having greater extensors CSA among women (Table 2). After adjusting for covariates, Tsimane women had 14.2 (95% CI: 9.6, 18.7) % greater extensors CSA than FHS women (Figure 2A; Table 4). Being younger, Tsimane, taller, and lighter were associated with having greater extensors density among women (Table 2). After adjusting for covariates, Tsimane women had 10.8 (4.1, 17.4) % higher extensors density than FHS women (Figure 2C; Table 4). The age-related difference in extensors CSA or density was similar between cohorts (Figure 2B, D; Table 5).

**Figure 2.**
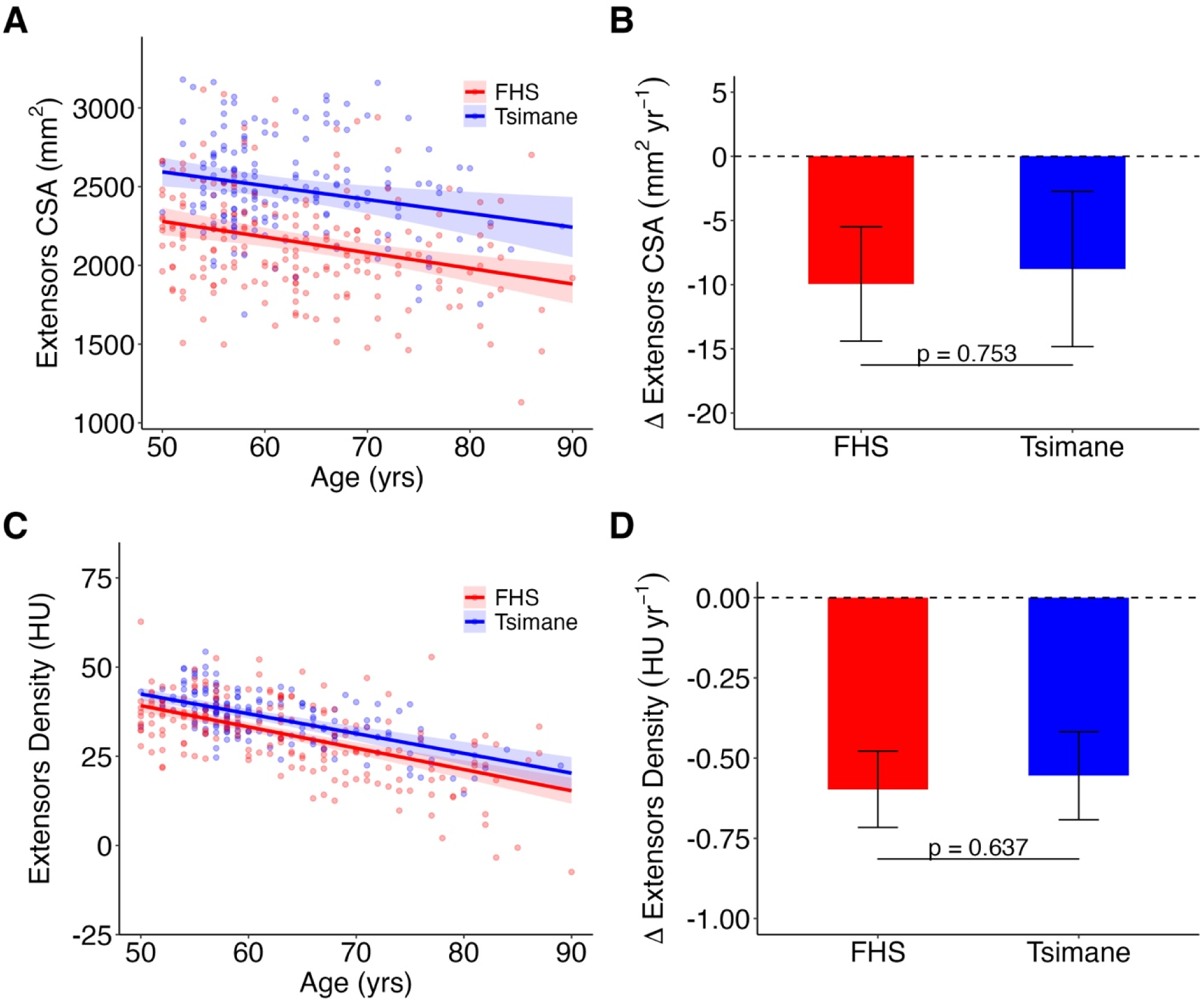
Tsimane women (*N*=165) had greater extensors cross-sectional area (CSA) and density than FHS women (*N*=205). Data points are values adjusted for height and weight. *p* values are for estimated marginal effects of age. HU, Hounsfield units. (A) Scatter plot and linear regression with 95% CI shading of extensors CSA versus age. (B) Bar plot of mean ± SD error bars of age-related differences in extensors CSA. (C) Scatter plot and linear regression with 95% CI shading of extensors density versus age. (D) Bar plot of mean ± SD error bars of age-related differences in extensors density.

**Table 2.** Linear regression models: effects of age, cohort, height, and weight on paraspinal muscle cross-sectional area (CSA) and density in women (Model 1). Model 2 includes an age–cohort interaction term. Model variables are not mean-centered.

| Parameter | Extensors CSA, mm<sup>2</sup> |  |  |  | Extensors Density, HU |  |  |  |
| --- | --- | --- | --- | --- | --- | --- | --- | --- |
|  | Model 1 |  | Model 2 |  | Model 1 |  | Model 2 |  |
|  | β (95% CI) | P | β (95% CI) | P | β (95% CI) | P | β (95% CI) | P |
| (Intercept) | 374 (-728, 1,477) | 0.505 | 375 (-730, 1,480) | 0.505 | 39.8 (15.8, 63.8) | **0.001** | 39.8 (15.6, 64.0) | **0.001** |
| Age (yrs) | -9.55 (-13.2, -5.87) | **<0.001** | -9.94 (-14.4, -5.49) | **<0.001** | -0.582 (-0.674, -0.490) | **<0.001** | -0.597 (-0.716, -0.478) | **<0.001** |
| Tsimane (ref: FHS) | 328 (222, 434) | **<0.001** | 256 (-204, 716) | 0.275 | 3.60 (1.38, 5.82) | **0.002** | 1.13 (-9.58, 11.8) | 0.836 |
| Height (cm) | 9.24 (2.89, 15.6) | **0.004** | 9.39 (2.96, 15.8) | **0.004** | 0.324 (0.186, 0.463) | **<0.001** | 0.330 (0.191, 0.469) | **<0.001** |
| Weight (kg) | 14.3 (11.8, 16.8) | **<0.001** | 14.3 (11.8, 16.8) | **<0.001** | -0.341 (-0.399, -0.283) | **<0.001** | -0.341 (-0.399, -0.283) | **<0.001** |
| Age–Tsimane | — | — | 1.17 (-6.10, 8.44) | 0.753 | — | — | 0.043 (-0.134, 0.219) | 0.637 |
| Adjusted R<sup>2</sup> | 0.382 |  | 0.380 |  | 0.506 |  | 0.502 |  |
| N | 370 |  | 370 |  | 370 |  | 370 |  |
| Parameter | Psoas major CSA, mm<sup>2</sup> |  |  |  | Psoas major Density, HU |  |  |  |
|  | Model 1 |  | Model 2 |  | Model 1 |  | Model 2 |  |
|  | β (95% CI) | P | β (95% CI) | P | β (95% CI) | P | β (95% CI) | P |
| (Intercept) | 184 (-183, 551) | 0.325 | 182 (-186, 550) | 0.331 | 32.2 (9.78, 54.5) | **0.005** | 32.4 (10.0, 54.8) | **0.005** |
| Age (yrs) | -1.73 (-2.96, -0.496) | **0.006** | -1.64 (-3.30, 0.018) | 0.053 | -0.230 (-0.308, -0.153) | **<0.001** | -0.246 (-0.338, -0.155) | **<0.001** |
| Tsimane (ref: FHS) | -143 (-178, -108) | **<0.001** | -132 (-278, 14.4) | 0.077 | -6.41 (-8.57, -4.25) | **<0.001** | -9.52 (-19.3, 0.266) | 0.057 |
| Height (cm) | 1.70 (-0.406, 3.81) | 0.113 | 1.68 (-0.458, 3.81) | 0.123 | 0.236 (0.107, 0.364) | **<0.001** | 0.240 (0.111, 0.369) | **<0.001** |
| Weight (kg) | 4.43 (3.57, 5.28) | **<0.001** | 4.43 (3.57, 5.28) | **<0.001** | -0.215 (-0.263, -0.167) | **<0.001** | -0.214 (-0.263, -0.166) | **<0.001** |
| Age–Tsimane | — | — | -0.173 (-2.48, 2.13) | 0.883 | — | — | 0.051 (-0.105, 0.207) | 0.521 |
| Adjusted R<sup>2</sup> | 0.647 |  | 0.646 |  | 0.329 |  | 0.328 |  |
| N | 370 |  | 370 |  | 370 |  | 370 |  |

HU, Hounsfield units; FHS, Framingham Heart Study

**Table 3.** Linear regression models: effects of age, cohort, height, and weight on paraspinal muscle cross-sectional area (CSA) and density in men (Model 1). Model 2 includes an age–cohort interaction term. Model variables are not mean-centered.

| Parameter | Extensors CSA, mm<sup>2</sup> |  |  |  | Extensors Density, HU |  |  |  |
| --- | --- | --- | --- | --- | --- | --- | --- | --- |
|  | Model 1 |  | Model 2 |  | Model 1 |  | Model 2 |  |
| | $\beta$ (95% CI) | P | $\beta$ (95% CI) | P | $\beta$ (95% CI) | P | $\beta$ (95% CI) | P |
| (Intercept) | 2,106 (683, 3,529) | **0.004** | 2,277 (836, 3,718) | **0.002** | 72.4 (45.4, 99.4) | **<0.001** | 77.6 (50.5, 105) | **<0.001** |
| Age (yrs) | -22.0 (-26.5, -17.5) | **<0.001** | -24.7 (-30.5, -18.9) | **<0.001** | -0.611 (-0.706, -0.516) | **<0.001** | -0.720 (-0.844, -0.597) | **<0.001** |
| Tsimane (ref: FHS) | -76.3 (-228, 75.6) | 0.324 | -481 (-1,050, 88.9) | 0.098 | 2.14 (-0.769, 5.05) | 0.149 | -12.6 (-24.0, -1.23) | **0.030** |
| Height (cm) | 5.44 (-2.53, 13.4) | 0.181 | 5.56 (-2.40, 13.5) | 0.171 | 0.132 (-0.020, 0.285) | 0.089 | 0.143 (-0.008, 0.294) | 0.063 |
| Weight (kg) | 19.0 (15.1, 22.9) | **<0.001** | 18.8 (14.9, 22.8) | **<0.001** | -0.238 (-0.313, -0.162) | **<0.001** | -0.244 (-0.318, -0.170) | **<0.001** |
| Age–Tsimane | — | — | 6.20 (-2.21, 14.6) | 0.148 | — | — | 0.246 (0.064, 0.428) | **0.008** |
| Adjusted R<sup>2</sup> | 0.579 |  | 0.596 |  | 0.414 |  | 0.417 |  |
| N | 371 |  | 371 |  | 371 |  | 371 |  |
| Parameter | Psoas major CSA, mm<sup>2</sup> |  |  |  | Psoas major Density, HU |  |  |  |
|  | Model 1 |  | Model 2 |  | Model 1 |  | Model 2 |  |
| | $\beta$ (95% CI) | P | $\beta$ (95% CI) | P | $\beta$ (95% CI) | P | $\beta$ (95% CI) | P |
| (Intercept) | 1,573 (925, 2,222) | **<0.001** | 1,612 (949, 2,275) | **<0.001** | 73.3 (53.0, 93.7) | **<0.001** | 74.2 (53.6, 94.8) | **<0.001** |
| Age (yrs) | -7.20 (-9.22, -5.18) | **<0.001** | -7.88 (-10.8, -4.99) | **<0.001** | -0.356 (-0.424, -0.288) | **<0.001** | -0.372 (-0.456, -0.288) | **<0.001** |
| Tsimane (ref: FHS) | -210 (-280, -139) | **<0.001** | -289 (-547, -29.9) | **0.029** | -3.35 (-5.53, -1.17) | **0.003** | -6.32 (-15.1, 2.44) | 0.157 |
| Height (cm) | -4.04 (-7.68, -0.396) | **0.030** | -3.98 (-7.63, -0.328) | **0.033** | 0.047 (-0.067, 0.162) | 0.417 | 0.048 (-0.067, 0.163) | 0.410 |
| Weight (kg) | 8.53 (6.64, 10.4) | **<0.001** | 8.47 (6.57, 10.4) | **<0.001** | -0.196 (-0.251, -0.141) | **<0.001** | -0.196 (-0.251, -0.141) | **<0.001** |
| Age–Tsimane | — | — | 1.20 (-2.58, 4.99) | 0.533 | — | — | 0.049 (-0.088, 0.186) | 0.486 |
| Adjusted R<sup>2</sup> | 0.593 |  | 0.590 |  | 0.282 |  | 0.280 |  |
| N | 371 |  | 371 |  | 371 |  | 371 |  |

HU, Hounsfield units; FHS, Framingham Heart Study

**Table 4.** Mean differences in paraspinal muscle cross-sectional area (CSA) and density with adjustment for age, height, and weight between Tsimane and Framingham Heart Study (FHS).

| Variable | Tsimane (N = 330)<br>Mean (95% CI) | FHS (N = 411)<br>Mean (95% CI) | % Diff (95% CI) |
| --- | --- | --- | --- |
| **Women** |  |  |  |
| Extensors |  |  |  |
| CSA, mm<sup>2</sup> | 2479.6 (2411.1, 2548.1) | 2151.8 (2096.1, 2207.5) | **14.2 (9.6, 18.7) \*\*\*** |
| Density, HU | 35.2 (33.8, 36.6) | 31.6 (30.3, 32.9) | **10.8 (4.1, 17.4) \*\*** |
| Psoas major |  |  |  |
| CSA | 487.5 (466.2, 508.7) | 630.1 (610.2, 649.9) | **-25.5 (-31.8, -19.3) \*\*\*** |
| Density | 34.0 (32.7, 35.4) | 40.4 (39.3, 41.6) | **-17.2 (-23.0, -11.4) \*\*\*** |
| **Men** |  |  |  |
| Extensors |  |  |  |
| CSA | 3023.5 (2929.9, 3117.1) | 3099.9 (3020.3, 3179.5) | -2.5 (-7.5, 2.5) |
| Density | 39.6 (37.9, 41.4) | 37.5 (35.9, 39.1) | 5.5 (-2.0, 13.1) |
| Psoas major |  |  |  |
| CSA | 885.8 (844.0, 927.6) | 1095.5 (1056.8, 1134.2) | **-21.2 (-28.3, -14.1) \*\*\*** |
| Density | 40.2 (38.8, 41.6) | 43.6 (42.5, 44.7) | **-8.0 (-13.2, -2.8) \*\*** |

HU, Hounsfield Units
\* $p < 0.05$ ; \*\* $p < 0.01$ ; \*\*\* $p < 0.001$ for estimated marginal means from linear regression
% Diff calculated as the difference between Tsimane and FHS means, divided by their average, and multiplied by 100

**Table 5.** Age-related differences in paraspinal muscle cross-sectional area (CSA) and density with adjustment for height and weight between Tsimane and Framingham Heart Study (FHS).

| Variable | Tsimane (N = 330) |  | FHS (N = 411) |  | % Diff, Mean Δ yr<sup>-1</sup> (95% CI) |
| --- | --- | --- | --- | --- | --- |
|  | Mean Δ yr<sup>-1</sup> (95% CI) | % Δ yr<sup>-1</sup> (95% CI) | Mean Δ yr<sup>-1</sup> (95% CI) | % Δ yr<sup>-1</sup> (95% CI) |  |
| Women |  |  |  |  |  |
| Extensors |  |  |  |  |  |
| CSA, mm<sup>2</sup> | -8.78 (-14.83, -2.73) | -0.34 (-0.57, -0.10) | -9.94 (-14.40, -5.49) | -0.44 (-0.63, -0.24) | -12.5 (65.2, -90.1) |
| Density, HU | -0.55 (-0.69, -0.42) | -1.30 (-1.62, -0.98) | -0.60 (-0.72, -0.48) | -1.53 (-1.83, -1.22) | -7.4 (23.3, -38.1) |
| Psoas major |  |  |  |  |  |
| CSA | -1.82 (-3.54, -0.10) | -0.36 (-0.69, -0.02) | -1.64 (-3.30, 0.02) | -0.25 (-0.51, 0.00) | 10.0 (143.5, -123.5) |
| Density | -0.20 (-0.33, -0.06) | -0.53 (-0.89, -0.17) | -0.25 (-0.34, -0.15) | -0.57 (-0.78, -0.36) | -23.1 (47.6, -93.8) |
| Men |  |  |  |  |  |
| Extensors |  |  |  |  |  |
| CSA | -18.51 (-25.03, -11.99) | -0.56 (-0.75, -0.36) | -24.71 (-30.48, -18.94) | -0.73 (-0.90, -0.56) | -28.7 (10.2, -67.6) |
| Density | -0.47 (-0.61, -0.34) | -0.99 (-1.28, -0.70) | -0.72 (-0.84, -0.60) | -1.57 (-1.84, -1.30) | **-41.2 (-10.7, -71.7) \*\*** |
| Psoas major |  |  |  |  |  |
| CSA | -6.68 (-9.34, -4.02) | -0.68 (-0.95, -0.41) | -7.88 (-10.76, -4.99) | -0.66 (-0.90, -0.42) | -16.5 (35.5, -68.5) |
| Density | -0.32 (-0.43, -0.21) | -0.72 (-0.97, -0.47) | -0.37 (-0.46, -0.29) | -0.77 (-0.94, -0.59) | -14.0 (25.5, -53.4) |

HU, Hounsfield Units
\* $p < 0.05$ ; \*\* $p < 0.01$ ; \*\*\* $p < 0.001$ for estimated marginal effects from linear regression
% Diff calculated as the difference between Tsimane and FHS mean $\Delta$ yr<sup>-1</sup>, divided by their average, and multiplied by 100

Linear regressions of age versus extensors CSA and density values unadjusted for covariates are found in Figure S5. On average, adjusting for covariates increased extensors CSA for Tsimane women by 216 mm^2^ and decreased extensors CSA for FHS women by 174 mm^2^ (Figure S5B). Adjusting for covariates decreased extensors density for Tsimane women by 1.2 HU and increased extensors density for FHS women by 1.0 HU (Figure S5D).

### Tsimane women had smaller and less-dense psoas major than FHS women

Being younger, FHS, and heavier were associated with having greater psoas major CSA among women (Table 2). After adjusting for covariates, Tsimane women had 25.5 (19.3, 31.8)% smaller psoas major CSA than FHS women (Figure 3A; Table 4). Being younger, FHS, taller, and lighter were associated with having greater psoas major density among women (Table 2). After adjusting for covariates, Tsimane women had 17.2 (11.4, 23.0) % lower psoas major density than FHS women (Figure 3C; Table 4). The age-related difference in psoas major CSA or density was similar between cohorts (Figure 3B, D; Table 5).

**Figure 3.**
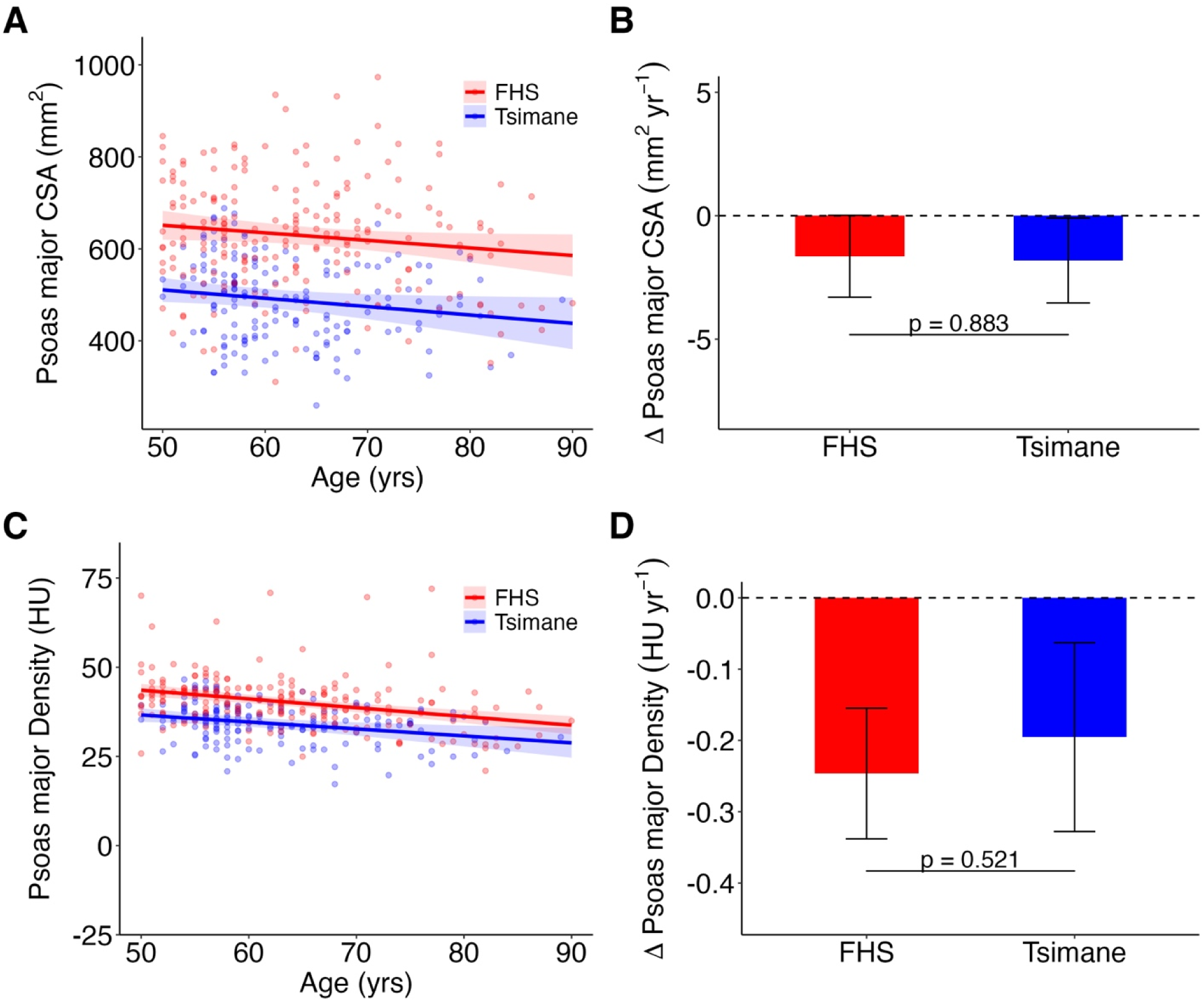
Tsimane women (*N*=165) had smaller psoas major cross-sectional area (CSA) and lower psoas major density than FHS women (*N*=205). Data points are values adjusted for height and weight. *p* values are for estimated marginal effects of age. HU, Hounsfield units. (A) Scatter plot and linear regression with 95% CI shading of psoas major CSA versus age. (B) Bar plot of mean ± SD error bars of age-related differences in psoas major CSA. (C) Scatter plot and linear regression with 95% CI shading of psoas major density versus age. (D) Bar plot of mean ± SD error bars of age-related differences in psoas major density.

Linear regressions of age versus psoas major CSA and density values unadjusted for covariates are found in Figure S6. On average, adjusting for covariates increased psoas major CSA for Tsimane women by 58 mm^2^ and decreased psoas major CSA for FHS women by 47 mm^2^ (Figure S6B). Adjusting for covariates decreased psoas major density for Tsimane women by 0.5 HU and increased psoas major density for FHS women by 0.4 HU (Figure S6D).

### Tsimane men had a smaller age-related difference in extensors density than FHS men

Being younger and heavier were associated with having greater extensors CSA and among men (Table 3). After adjusting for covariates, Tsimane and FHS men had similar extensors CSA (Figure 4A; Table 4). Being younger and lighter were associated with having greater extensors density among men (Table 3). After adjusting for covariates, Tsimane and FHS men had similar extensors density (Figure 4C; Table 4). For men, age interacts with cohort, along with being lighter, to influence extensors density. After adjusting for covariates, the age-related difference in extensors density was 41.2 (10.7, 71.7) % smaller for Tsimane men than FHS men (Figure 4D; Table 5).

**Figure 4.**
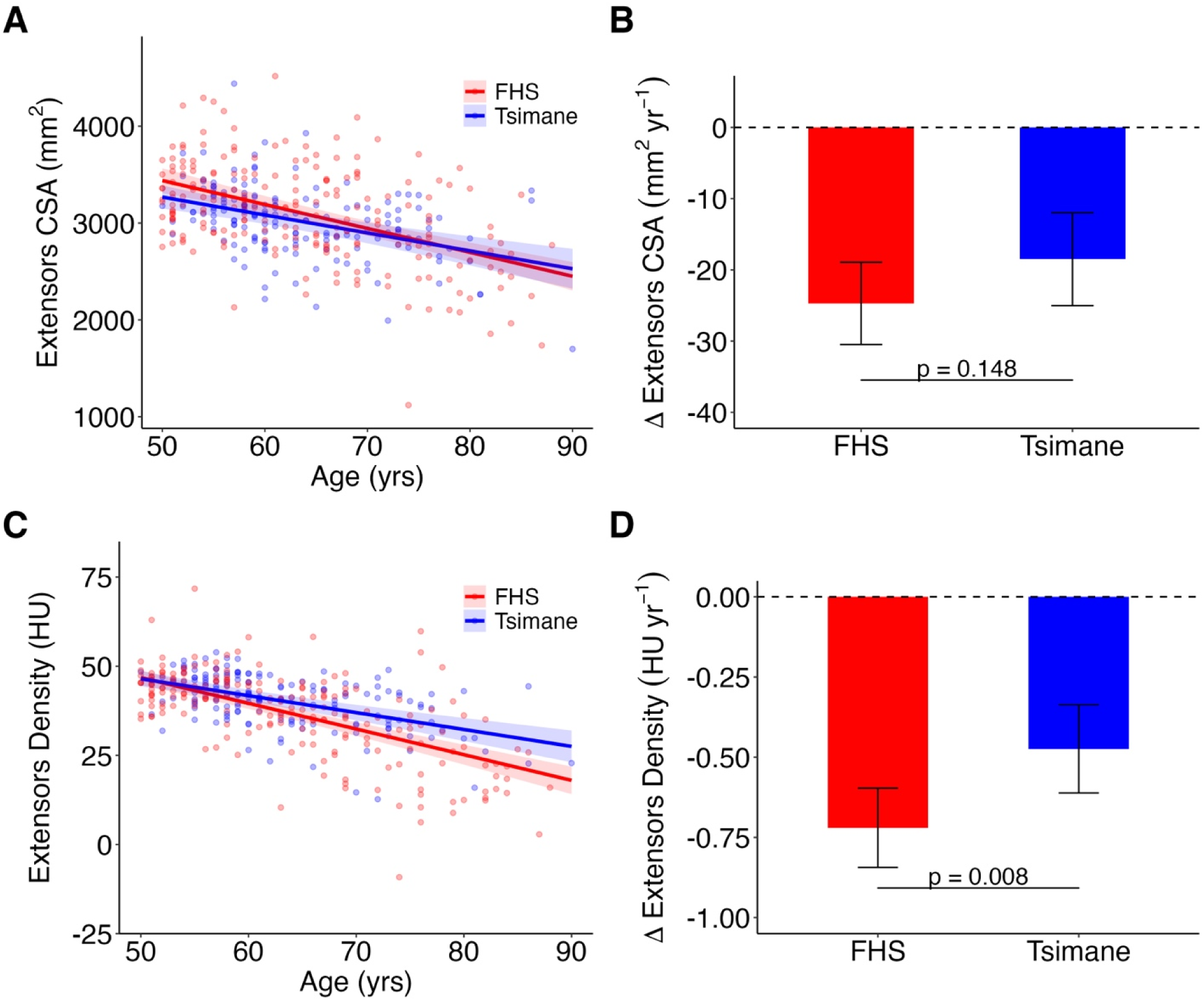
Tsimane men (*N*=165) had a smaller age-related difference in extensors density than FHS men (*N*=206). Data points are values adjusted for height and weight. *p* values are for estimated marginal effects of age. HU, Hounsfield units. (A) Scatter plot and linear regression with 95% CI shading of extensors CSA versus age. (B) Bar plot of mean ± SD error bars of age-related differences in extensors CSA. (C) Scatter plot and linear regression with 95% CI shading of extensors density versus age. (D) Bar plot of mean ± SD error bars of age-related differences in extensors density.

Linear regressions of age versus extensors CSA and density values unadjusted for covariates are found in Figure S7. On average, adjusting for covariates increased extensors CSA for Tsimane men by 340 mm^2^ and decreased extensors CSA for FHS men by 272 mm^2^ (Figure S7B). Adjusting for covariates decreased extensors density for Tsimane men by 2.5 HU and increased extensors density for FHS men by 2.0 HU (Figure S7D).

### Tsimane men had smaller and less-dense psoas major than FHS men

Being younger, FHS, shorter, and heavier were associated with having greater psoas major CSA among men (Table 3). After adjusting for covariates, Tsimane men had 21.2 (14.1, 28.3) % smaller psoas major CSA than FHS men (Figure 5A; Table 4). Being younger, FHS, and lighter were associated with having greater psoas major density among men (Table 3). After adjusting for covariates, Tsimane men had 8.0 (2.8, 13.2) % lower psoas major density than FHS men (Figure 5C; Table 4). The age-related difference in psoas major CSA or density was similar between cohorts (Figure 5B, D; Table 5).

**Figure 5.**
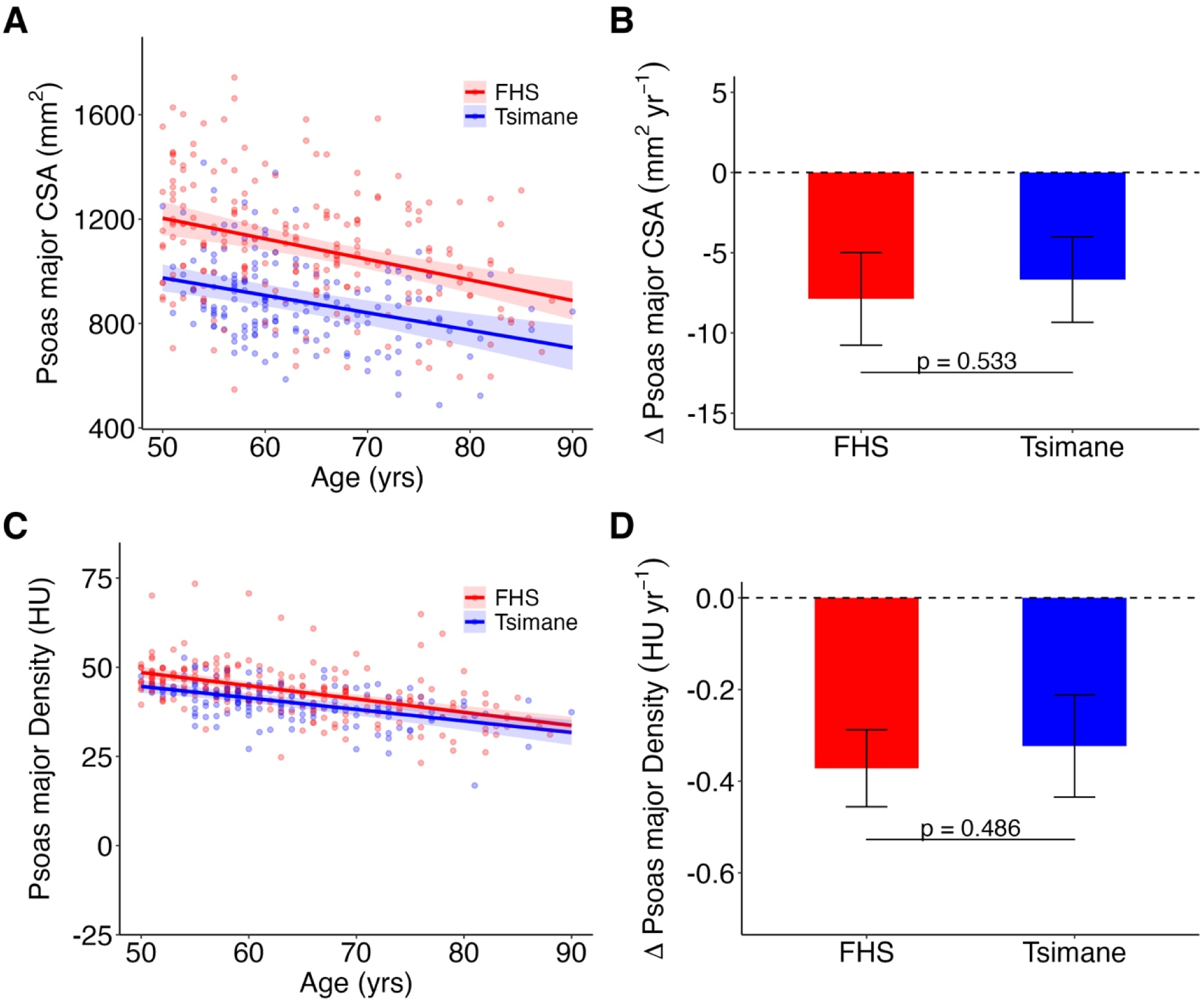
Tsimane men (*N*=165) had smaller psoas major cross-sectional area (CSA) and lower psoas major density than FHS men (*N*=206). Data points are values adjusted for height and weight. *p* values are for estimated marginal effects of age. HU, Hounsfield units. (A) Scatter plot and linear regression with 95% CI shading of psoas major CSA versus age. (B) Bar plot of mean ± SD error bars of age-related differences in psoas major CSA. (C) Scatter plot and linear regression with 95% CI shading of psoas major density versus age. (D) Bar plot of mean ± SD error bars of age-related differences in psoas major density.

Linear regressions of age versus psoas major CSA and density values unadjusted for covariates are found in Figure S8. On average, adjusting for covariates increased psoas major CSA for Tsimane men by 96 mm^2^ and decreased psoas major CSA for FHS men by 77 mm^2^ (Figure S6B). Adjusting for covariates decreased psoas major density for Tsimane men by 2.6 HU and increased psoas major density for FHS men by 2.1 HU (Figure S6D).

## Discussion

Our results show that adults from populations at opposite ends of the industrialization spectrum have a different balance in the size and density of trunk extensors and psoas major. Adjusting for age, height, and weight, Tsimane women had greater extensors size and density, but lower psoas major size and density, compared to FHS women. Tsimane men had a smaller age-related difference in extensors density, but lower psoas major size and density, than FHS men. These results provide mixed evidence for our hypotheses. We partially confirmed our prediction that Tsimane extensors size and density are higher than FHS, with smaller age-related differences, likely because Tsimane engage in more physical activity beginning early in life that is maintained throughout older age. The finding that Tsimane have lower covariate-adjusted psoas major size and density could be explained by engagement in different types of physical activity, or by Tsimane experiencing high energy expenditure (i.e., from high physical activity and pathogen burden), which could potentially limit growth and maintenance of muscle tissue. Future work is required, however, to test how exposome features (i.e., physical activity, pathogen burden, diet, reproduction) influence the size and density of trunk extensors and psoas major.

The FHS results are consistent with prior work that showed paraspinal muscle size and density decrease with advancing age [27,58,59,64,65]. Deficits in paraspinal muscle morphology are associated with back pain and vertebral fractures [10–13]. Our observation that Tsimane women have higher covariate-adjusted extensors size and density would suggest they may be better protected from vertebral fractures than FHS women. Conversely, lower psoas major morphology among Tsimane than FHS might suggest Tsimane are at higher risk of vertebral fracture. Prior work showed that prevalent thoracic vertebral fractures are twice as common among Tsimane women than US women (and three times as common for Tsimane men vs. US men) [66]. In that study, lower thoracic vertebral bone mineral density (BMD) was associated with prevalent vertebral fractures for Tsimane women, but not men. For Tsimane men, habitual and intense subsistence work, hence potentially high vertebral loading, was cited as a potential explanation for their prevalent thoracic vertebral fractures [67]. Higher vertebral loading contributes to higher load-to-strength ratios in the spine, potentially increasing fracture risk [68,69]. Higher thoracic vertebral fracture prevalence among Tsimane, combined with our finding of differences in trunk extensor and psoas major morphology between Tsimane and FHS, suggests associations between paraspinal muscles and vertebral fractures may be more intricate.

Cohort differences in paraspinal muscle size and density may be related to different exposomes (e.g., physical activity, diet) between cohorts. Trunk extensor size can be increased by exercise therapies that focus on motor control, stabilization, and muscle strengthening [48] and physical activity is positively associated with paraspinal muscle morphology and skeletal muscle mass [49,70]. Tsimane engage in more physical activity than FHS and perform subsistence-oriented tasks, like collecting water and firewood, that are rare among FHS; Tsimane also frequently pick up and carry children with the hands [40,43,44,71]. Living in the Amazon basin, Tsimane also traverse hills, muddy embankments, and forested terrains on foot; this requires they walk with more flexed trunks, hips, and knees [72]. Such activities and movements involving forward bending, lifting, and joint flexion require high levels of trunk extensors neuromuscular activity; over time, these muscle activation patterns may increase extensors size and density [22,73,74]. In addition, higher visceral adipose tissue is associated with lower paraspinal muscle density [75]. The Tsimane diet is low in fat compared to FHS [75–77] and Tsimane have relatively low amounts of ectopic adipose tissue [78]. Therefore, we speculate that different physical activity and diet patterns may explain the greater covariate-adjusted size and density of extensor muscles we observed for Tsimane women.

Physical activity and diet, however, would not appear to explain the smaller size and density of Tsimane psoas major muscles. This apparent contradiction may be due to psoas major’s function during sitting. Psoas major flexes the trunk and hip and laterally stabilizes the anterior lumbar spine; it is highly active during sitting with a straight-back posture [9,79]. We speculate that cohort differences in sitting posture may explain the relatively smaller and less-dense psoas major of Tsimane. Chairs are uncommon in subsistence-oriented societies; instead, people squat, kneel, sit on the ground, and adopt various non-chair sitting postures [80,81]. Such postures may involve more curved, rather than straight, backs [82]. Chair-based sitting, which is ubiquitous in post-industrialized societies like the US, may involve more straight-back postures [83–85]. Repeated-measures studies are required to test whether sitting posture influences paraspinal muscle morphology independent of other exposome features.

Paraspinal muscle size and density may also affect spinal curvatures. Having larger lumbar erector spinae is associated with having higher lumbar lordosis, the sagittal convex curvature in the lower back [86]. On average, women have higher lumbar lordosis than men [87]. This sex difference has been interpreted as an adaptation related to pregnancy: a higher lordosis shifts the weight of the fetus more posteriorly during pregnancy, closer to the maternal hip and lumbosacral joint centers; this shift reduces the flexor torque generated by the weight of the fetus and, consequently, reduces the extensor torque the mother’s trunk and hip extensor muscles must generate to maintain upright posture [23]. We speculate that the larger and denser trunk extensors we observed for Tsimane women may be partly related to the high average total fertility rate of Tsimane (9 live births per woman [88,89]). Total fertility rate is unavailable for FHS, but it is undoubtedly lower than for Tsimane; in a study of 2,357 FHS women, the mean number of pregnancies was 2.3 [90]. A higher total fertility rate may be associated with changes to spinal curvatures and paraspinal muscle morphology. Also, as mentioned earlier, Tsimane frequently pick up and carry children with the hands. A high total fertility rate likely means this behavior is performed often, and for many years, among both women and men.

### Limitations

This study’s sample of hundreds of participants aged 50–90 years from two population-based cohorts at opposite ends of the industrialization spectrum was observational and cross-sectional, and thus we are unable to assess causality. Although we adjusted for cohort differences in body size on paraspinal muscle CSA and density, it was beyond the scope of this study to determine the effect of specific exposome features on muscle morphology. Future work should identify how much variation in paraspinal muscle CSA and density is explained by exposome features, including physical activity, diet, reproduction, and pathogen burden. We measured paraspinal muscle morphology in the lumbar spine region. Prior work has shown that thoracic paraspinal muscle morphology is associated with prevalent thoracic vertebral fractures and thoracic kyphosis [91–93]. Future work should determine cohort differences in thoracic paraspinal muscle morphology as well as prevalent vertebral fractures and other spine pathologies (e.g., hyperkyphosis, disc degeneration). Non-paraspinal trunk muscles (e.g., rectus abdominis, latissimus dorsi, the obliques) indirectly support and move the spine during postural and locomotor behaviors. Future work should determine age-related patterns in the size and density of non-paraspinal trunk muscles between post-industrialized and non-industrialized population-based cohorts.

## Conclusion

After adjusting for body size, non-industrialized Tsimane forager-horticulturalist women had larger and denser trunk extensor muscles compared to post-industrialized FHS women. In contrast, Tsimane had smaller and less-dense psoas major muscles than FHS. Tsimane men had a smaller age-related difference in extensors density than FHS. Other than this, we did not observe age-related differences in paraspinal muscle morphology between cohorts. These results show that adults from populations at opposite ends of the industrialization spectrum have a different balance in the size and density of trunk extensors versus psoas major muscles. Paraspinal muscle size and density have previously been linked to back pain and vertebral fractures in post-industrialized populations. Future work should determine these associations in non-industrialized populations, like the Tsimane. Future work should also identify the influence of exposome features, like physical activity and diet, on paraspinal muscle size and density in populations across the industrialization spectrum. Such knowledge would improve our understanding of how to mitigate age-related declines in paraspinal muscle health.

## Acknowledgements

The authors thank the Tsimane and Framingham communities for their participation. We thank the THLHP and FHS teams for collecting data. We thank Elaine Yu, Adele Carty, Jane Wang, Crystabella Nevarez, and Keity Okazaki for helpful discussions pertaining to the manuscript. We extend additional appreciation to funding sources including the US National Institutes of Health/National Institute Of Diabetes And Digestive And Kidney Diseases (T32-DK007028); US National Institutes of Health/National Institute of Arthritis, Musculoskeletal and Skin Diseases (R01-AR073019); US National Institutes of Health/National Heart, Lung, and Blood Institute (NHLBI) Framingham Heart Study (NIH/NHLBI Contract N01-HC-25195); French National Research Agency (ANR) under the Investments for the Future (Investissements d’Avenir) program (ANR-17-EURE-0010 to the Toulouse School of Economics); US National Institutes of Health/National Institute on Aging (R01-AG054442 and R01-AG024119); US National Science Foundation (BCS1748282); Center for Evolutionary Medicine at Arizona State University; and University of California Santa Barbara Academic Senate. Funding sources had no role in research conduct, study design or article preparation.

## Data Availability Statement

All data and code needed to evaluate and reproduce the results in the paper are available from the authors upon reasonable request. This study did not generate new materials. For THLHP, individual-level data are stored in the THLHP Data Repository and are available through restricted access for ethical reasons. THLHP’s highest priority is the safeguarding of human subjects and minimization of risk to study participants. The THLHP adheres to the CARE Principles for Indigenous Data Governance, which assure that the Tsimane 1) have sovereignty over how data are shared, 2) are the primary gatekeepers determining ethical use, 3) are actively engaged in the data generation and 4) derive benefit from data generated and shared use whenever possible. The THLHP is also committed to the FAIR Principles to facilitate data use. Requests for individual-level data should take the form of an application that minimally details the exact uses of the data and the research questions to be addressed, procedures that will be employed for data security and individual privacy, potential benefits to the study community, and procedures for assessing and minimizing stigmatizing interpretations of the research results (see the following webpage for links to the data sharing policy and data request forms: https://tsimane.anth.ucsb.edu/data.html). Requests for individual-level data will require institutional IRB approval (even if exempt) and will be reviewed by an Advisory Council composed of tribal leaders, tribal community members, Bolivian scientists, and the THLHP leadership. The study authors and the Tsimane leadership are committed to open science and are available to assist interested investigators in preparing data access requests.

