## Supplemental Information for "Comparison of lumbar paraspinal muscle size and density between members of the US Framingham Heart Study and Tsimane forager-horticulturalists of Bolivia"

**Comparing manual vs. automated measurements of muscle morphology**

To evaluate the agreement between manual and automated measurements of muscle cross-sectional area (CSA) and density, we used linear regression analysis and Bland-Altman plots. In linear regression, for Framingham Heart Study (FHS, *N* = 272), *R^2^* values were between 0.88–0.94 and linear regression slopes were between 0.94–1.00 (Figure S1A & C, Figure S2A & C). For Tsimane (*N* = 40), values for *R^2^* were between 0.89–0.98 and linear regression slopes were between 0.90–1.00 (Figure S3A & C, Figure S4A & C).

In Bland-Altman plots, there was no evidence of systemic bias (all 95% CI intervals included 0). For FHS, mean [95% CI] bias for extensors CSA was 122 [-240, 483] mm^2^ (Figure S1B) and -14 [-248, 221] mm^2^ for psoas major CSA (Figure S2B). Mean bias for extensors density was -1.2 [-7.4, 4.5] Hounsfield units (HU) (Figure S1D) and -1.4 [-7.5, 4.7] HU for psoas major density (Figure S2D). For Tsimane, mean bias for extensors CSA was 163 [-7, 333] mm^2^ (Figure S3B) and 2 [-82, 86] mm^2^ for psoas major CSA (Figure S4B). Mean bias for extensors density was -1.2 [-3.4, 1.1] HU (Figure S3D) and -0.2 [-5.0, 4.6] HU for psoas major density (Figure S4D).

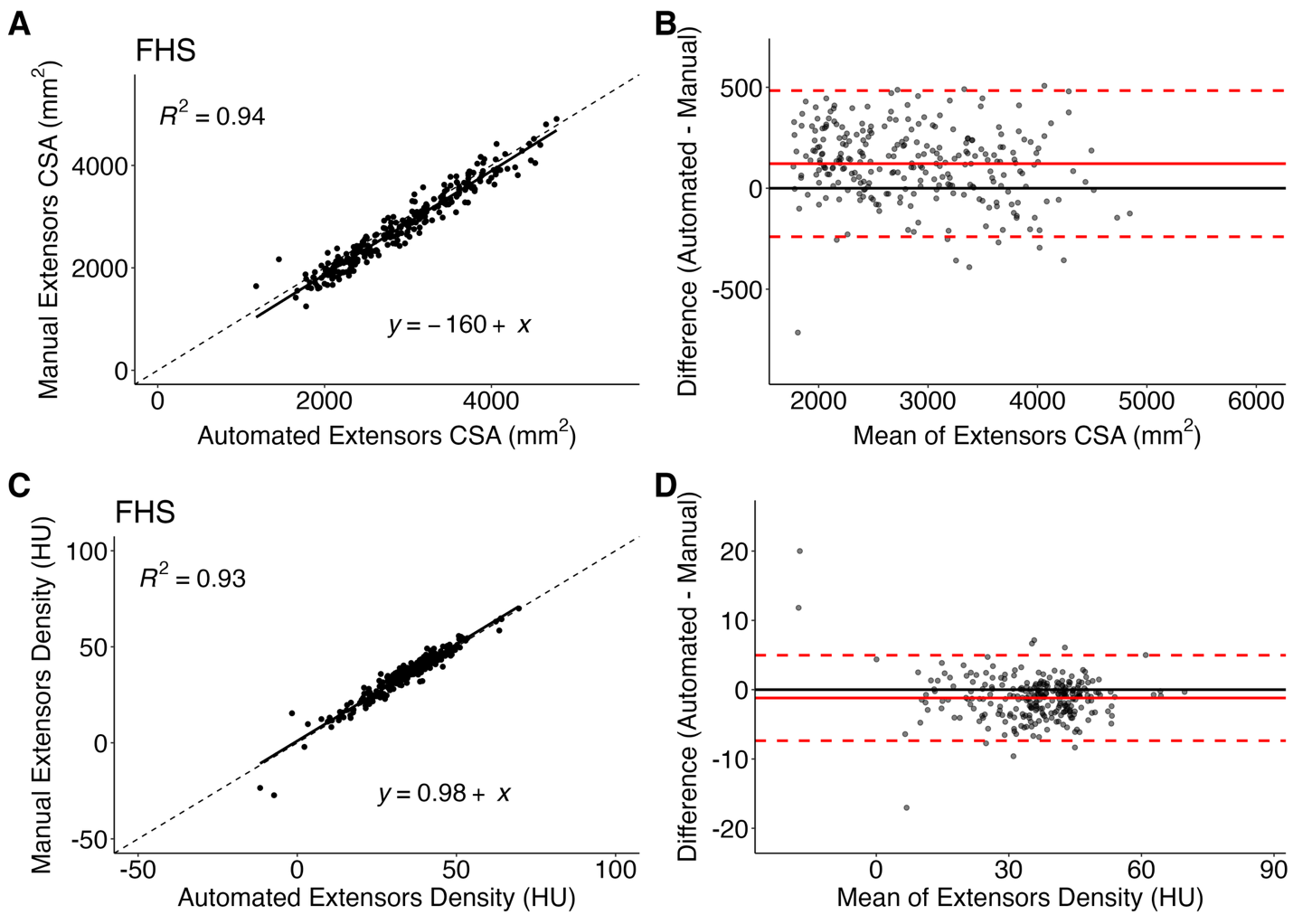

**Figure S1.** Scatter plots and linear regressions and Bland-Altman plots of manual versus automated measurements of extensors cross-sectional area (CSA) and density for FHS (*N* = 272). HU, Hounsfield units. (A–B) Extensors CSA. (C–D) Extensors density.

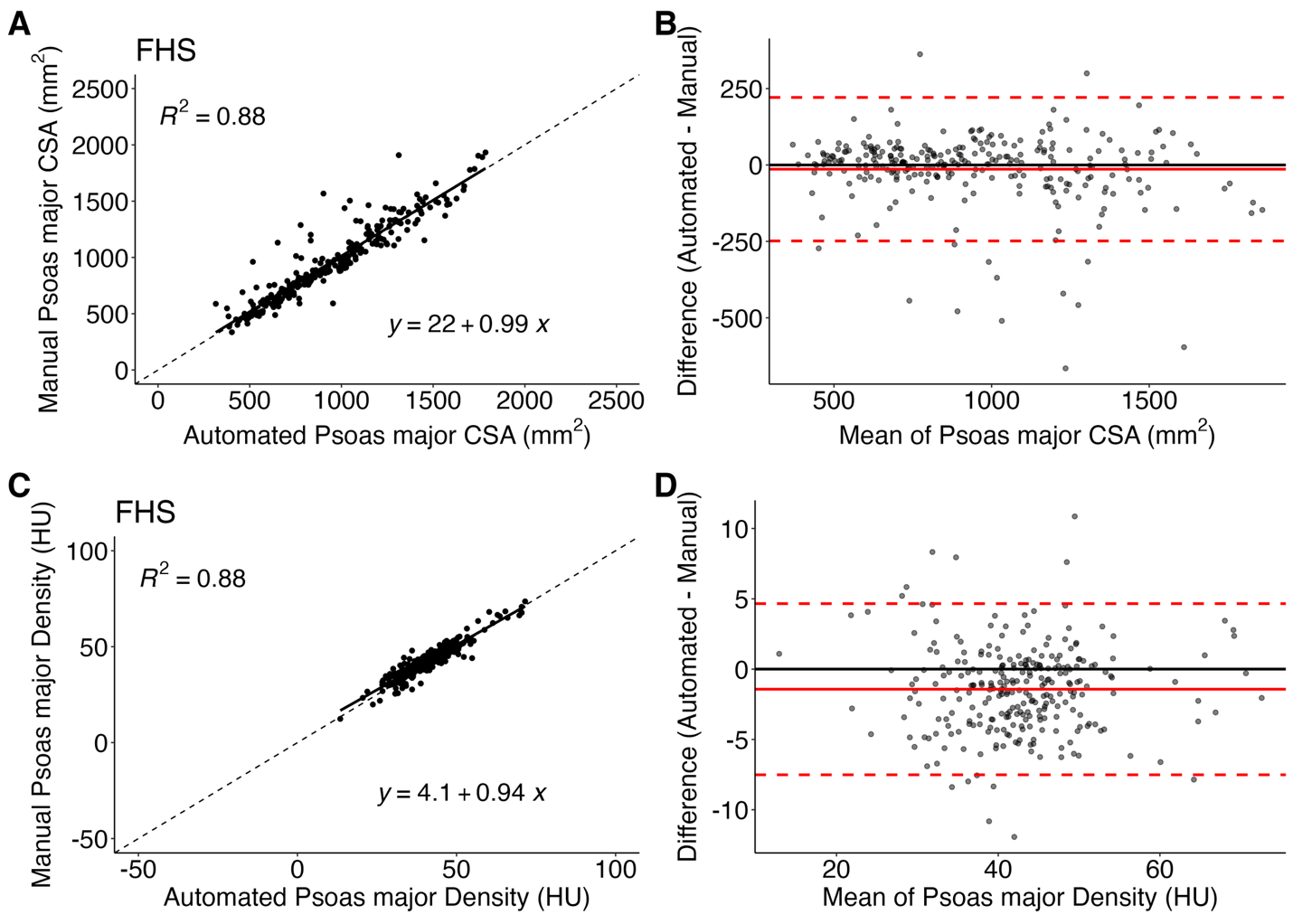

**Figure S2.** Scatter plots and linear regressions and Bland-Altman plots of manual versus automated measurements of psoas major cross-sectional area (CSA) and density for FHS (*N* = 272). HU, Hounsfield units. (A–B) Psoas major CSA. (C–D) Psoas major density.

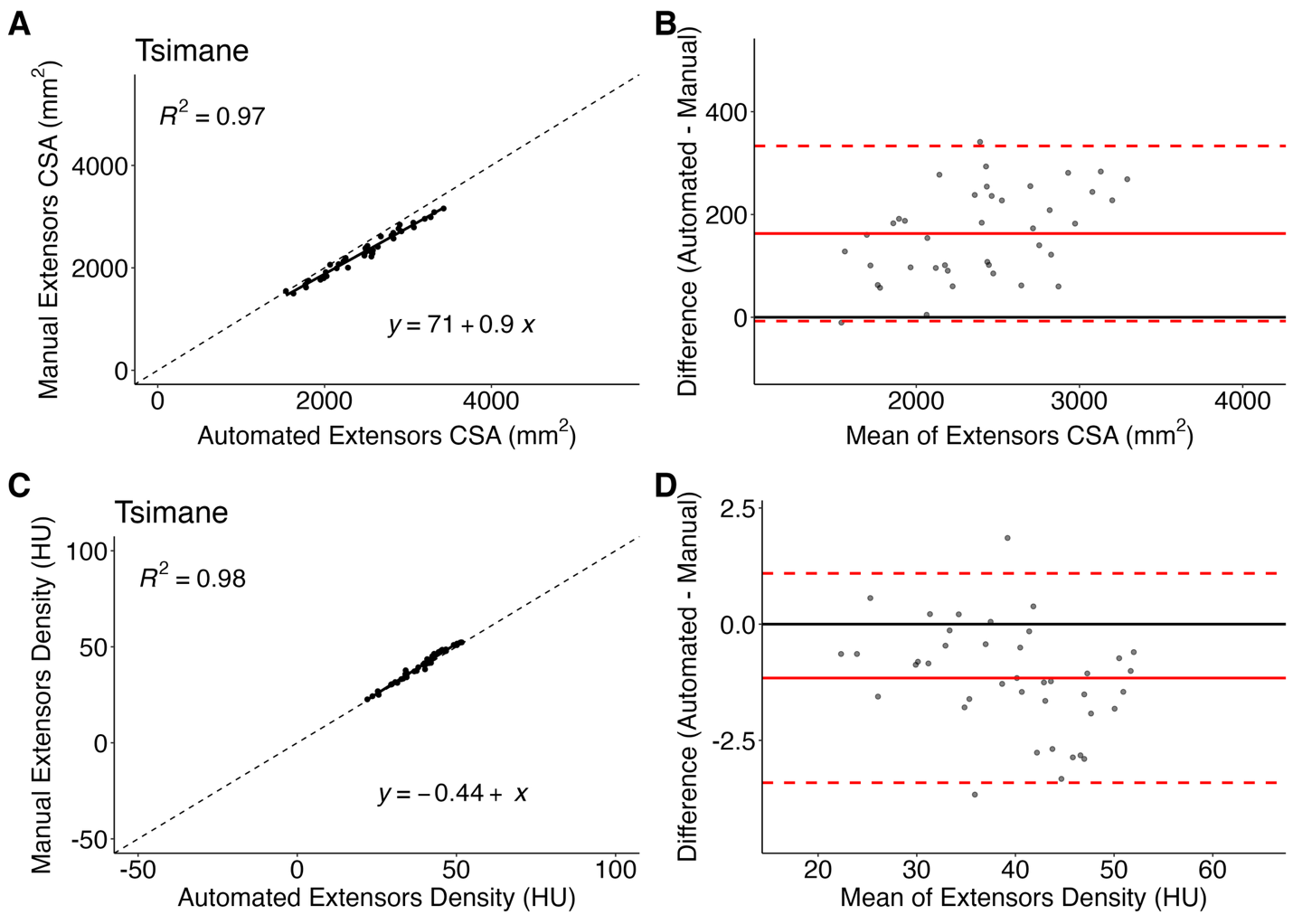

**Figure S3.** Scatter plots and linear regressions and Bland-Altman plots of manual versus automated measurements of extensors cross-sectional area (CSA) and density for Tsimane (*N* = 40). HU, Hounsfield units. (A–B) Extensors CSA. (C–D) Extensors density.

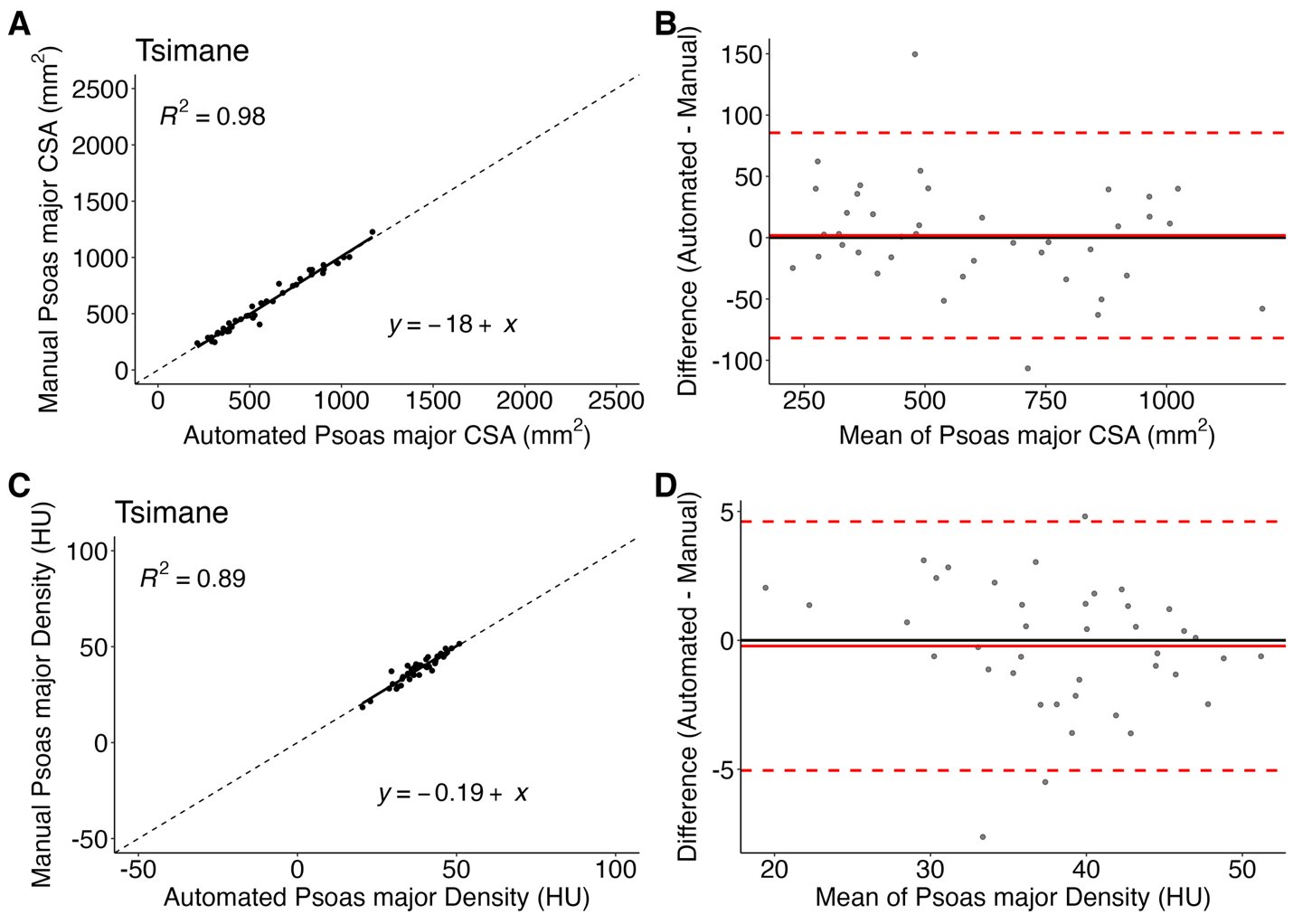

**Figure S4.** Scatter plots and linear regressions and Bland-Altman plots of manual versus automated measurements of psoas major cross-sectional area (CSA) and density for Tsimane (*N* = 40). HU, Hounsfield units. (A–B) Psoas major CSA. (C–D) Psoas major density.

**Table S1**. Descriptive statistics of paraspinal muscle cross-sectional area (CSA) and density values unadjusted for covariates for Tsimane and Framingham Heart Study (FHS).

| **Variable** | ***N*** | **Mean** | **SD** | **Min** | **Max** |
| --- | --- | --- | --- | --- | --- |
| CSA, mm^2^ |  |  |  |  |  |
| Tsimane Women |  |  |  |  |  |
| Extensors | 165 | 2273.8 | 391.4 | 1248.2 | 3292.9 |
| Psoas Major | 165 | 429.8 | 102.9 | 180.8 | 759.1 |
| Tsimane Men |  |  |  |  |  |
| Extensors | 165 | 2686.8 | 437.4 | 1222.3 | 4693.2 |
| Psoas Major | 165 | 793.2 | 184.5 | 329.3 | 1329.8 |
| FHS Women |  |  |  |  |  |
| Extensors | 205 | 2317.3 | 409.7 | 1452.5 | 3673.0 |
| Psoas Major | 205 | 676.5 | 143.4 | 354.0 | 1098.5 |
| FHS Men |  |  |  |  |  |
| Extensors | 206 | 3369.6 | 606.2 | 1179.5 | 5168.5 |
| Psoas Major | 206 | 1168.8 | 256.1 | 674.5 | 2133.0 |
| Density, HU |  |  |  |  |  |
| Tsimane Women |  |  |  |  |  |
| Extensors | 165 | 36.7 | 8.1 | 13.6 | 57.5 |
| Psoas Major | 165 | 34.7 | 5.6 | 17.7 | 50.7 |
| Tsimane Men |  |  |  |  |  |
| Extensors | 165 | 42.7 | 7.7 | 14.7 | 57.2 |
| Psoas Major | 165 | 43.0 | 5.1 | 20.9 | 55.3 |
| FHS Women |  |  |  |  |  |
| Extensors | 205 | 30.4 | 11.8 | -7.3 | 62.4 |
| Psoas Major | 205 | 39.9 | 8.4 | 13.5 | 70.5 |
| FHS Men |  |  |  |  |  |
| Extensors | 206 | 34.8 | 12.3 | -11.7 | 69.6 |
| Psoas Major | 206 | 41.3 | 8.0 | 22.0 | 71.6 |

HU, Hounsfield units

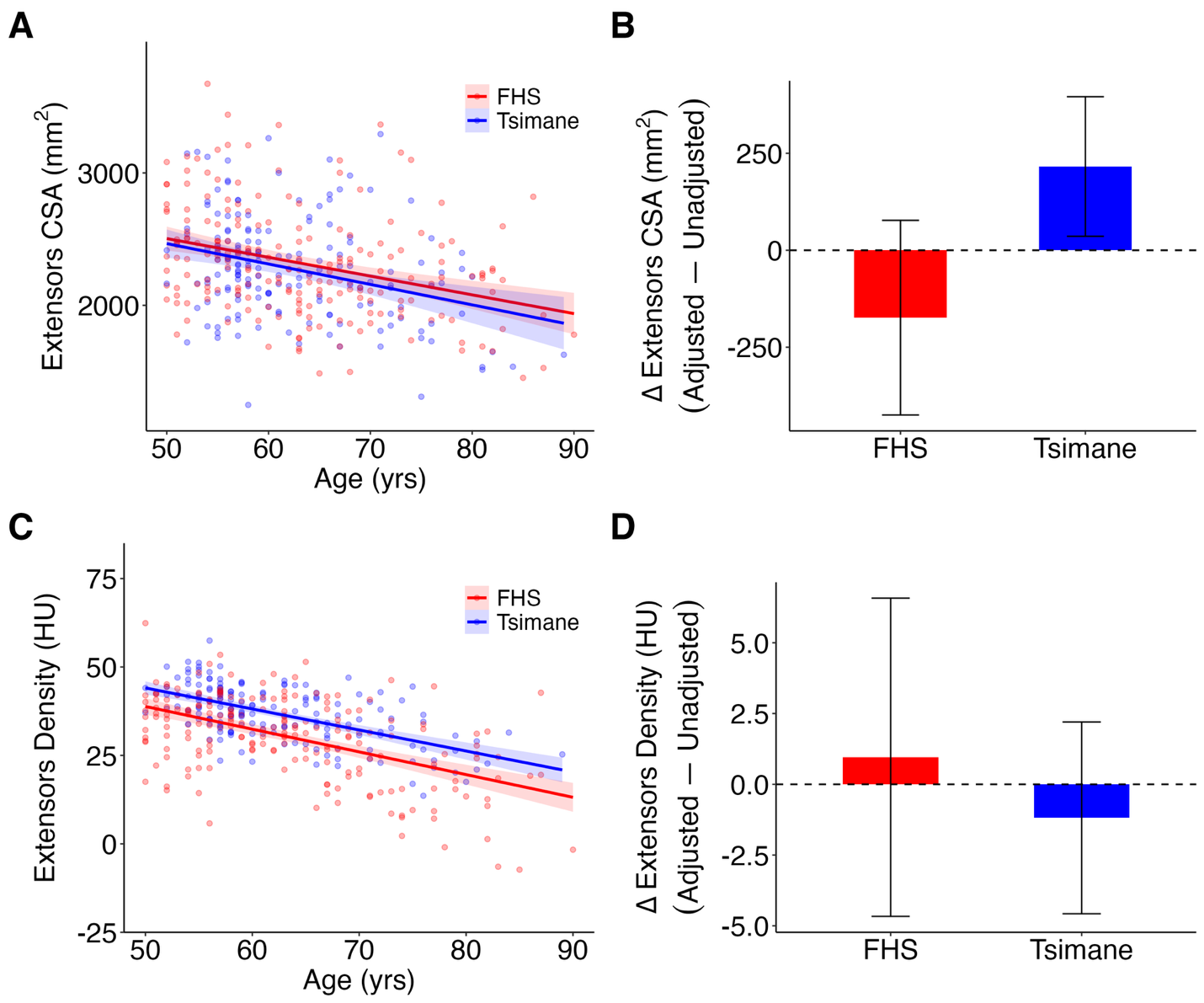

**Figure S5.** Extensors cross-sectional area (CSA) and density values unadjusted for height and weight for women, and the effect of adjusting for height and weight on extensors CSA and density. HU, Hounsfield units. (A) Scatter plot and linear regression with 95% CI shading of unadjusted extensors CSA with age. (B) Bar plot of mean ± SD error bars for adjusted–unadjusted extensors CSA. (C) Scatter plot and linear regression with 95% CI shading of unadjusted extensors density with age. (D) Bar plot of mean ± SD error bars for adjusted–unadjusted extensors density.

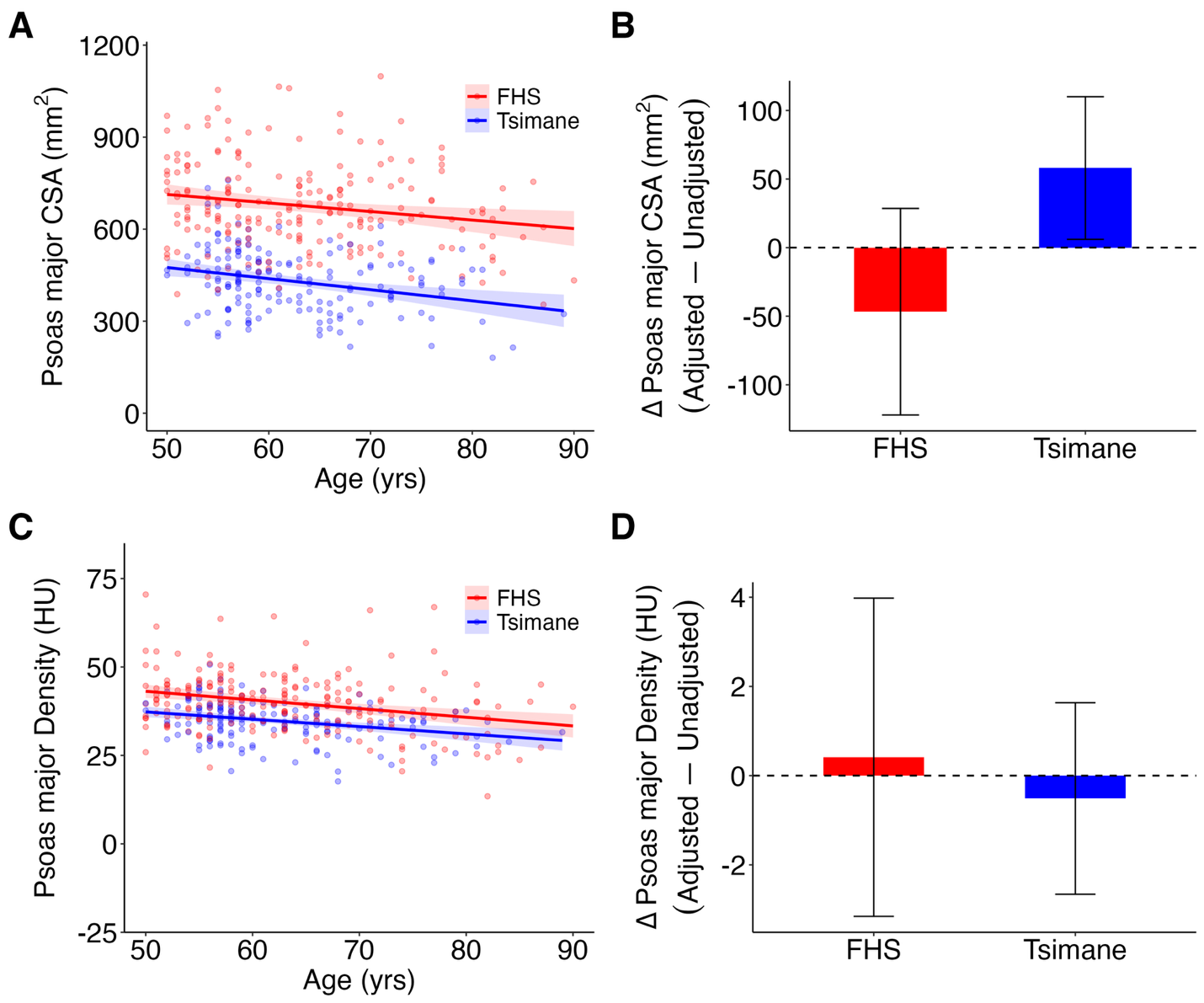

**Figure S6.** Psoas major cross-sectional area (CSA) and density values unadjusted for height and weight for women, and the effect of adjusting for height and weight on psoas major CSA and density. HU, Hounsfield units. (A) Scatter plot and linear regression with 95% CI shading of unadjusted psoas major CSA with age. (B) Bar plot of mean ± SD error bars for adjusted–unadjusted psoas major CSA. (C) Scatter plot and linear regression with 95% CI shading of unadjusted psoas major density with age. (D) Bar plot of mean ± SD error bars for adjusted–unadjusted psoas major density.

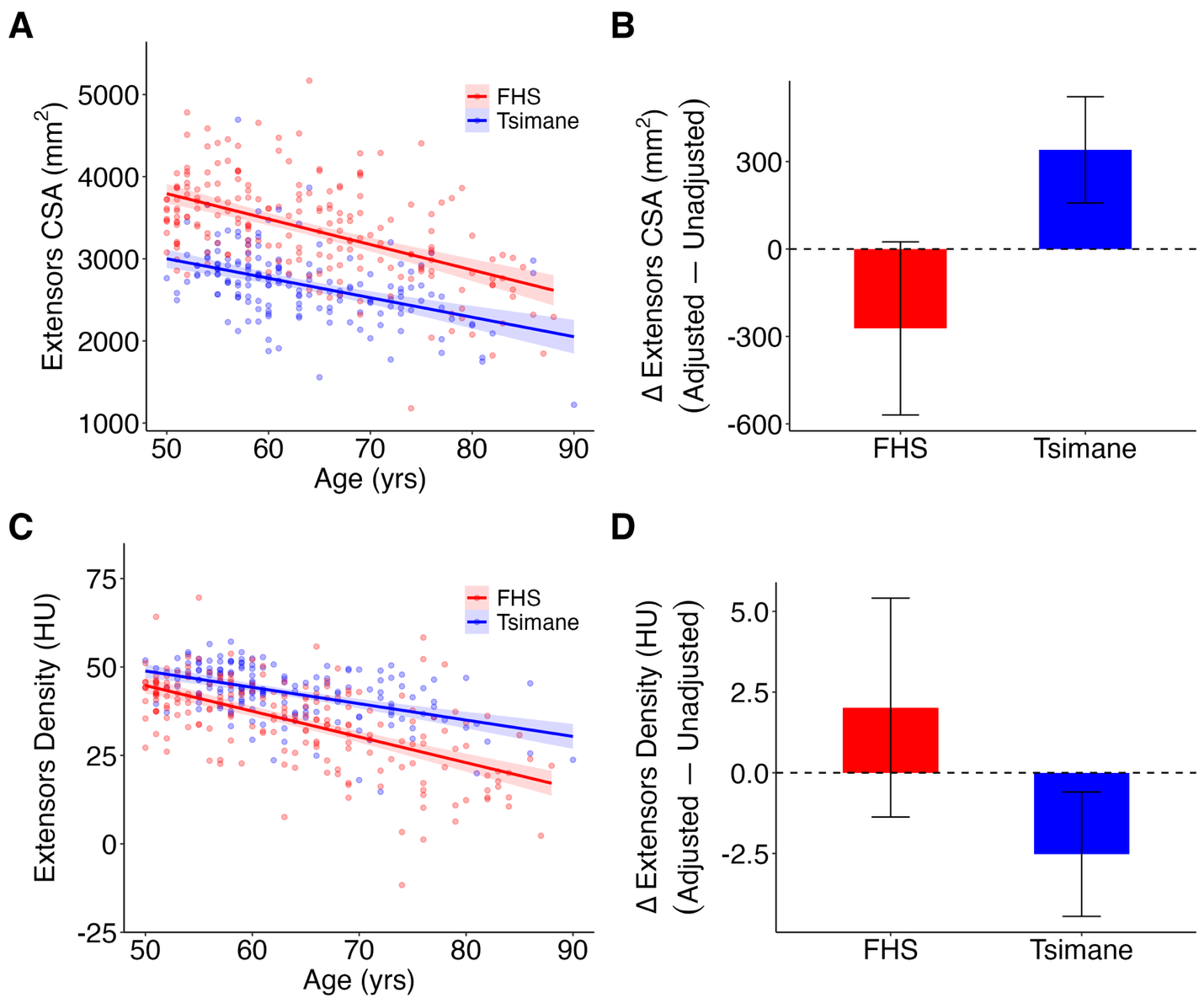

**Figure S7.** Extensors cross-sectional area (CSA) and density values unadjusted for height and weight for men, and the effect of adjusting for height and weight on extensors CSA and density. HU, Hounsfield units. (A) Scatter plot and linear regression with 95% CI shading of unadjusted extensors CSA with age. (B) Bar plot of mean ± SD error bars for adjusted–unadjusted extensors CSA. (C) Scatter plot and linear regression with 95% CI shading of unadjusted extensors density with age. (D) Bar plot of mean ± SD error bars for adjusted–unadjusted extensors density.

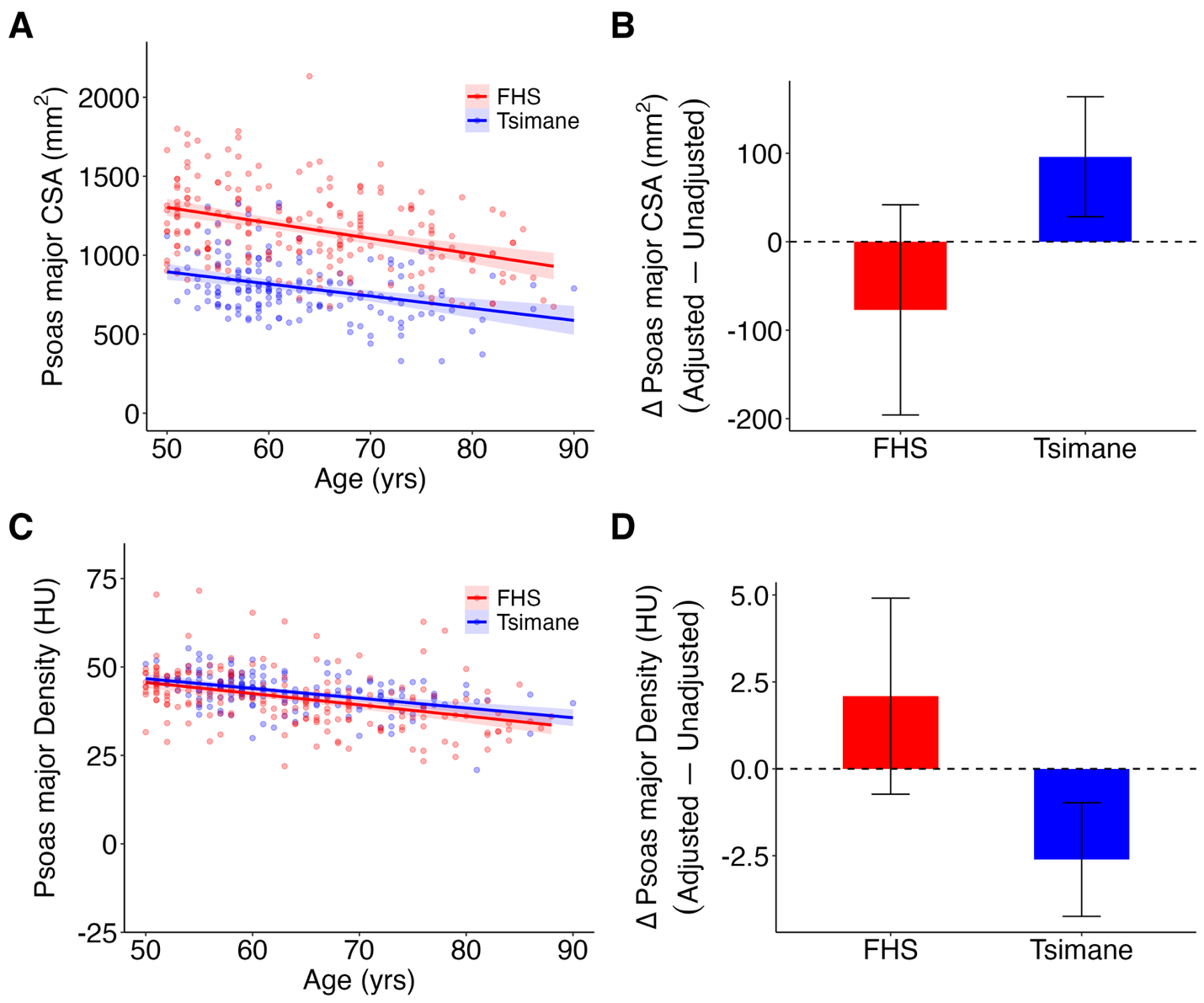

**Figure S8.** Psoas major cross-sectional area (CSA) and density values unadjusted for height and weight for men, and the effect of adjusting for height and weight on psoas major CSA and density. HU, Hounsfield units. (A) Scatter plot and linear regression with 95% CI shading of unadjusted psoas major CSA with age. (B) Bar plot of mean ± SD error bars for adjusted–unadjusted psoas major CSA. (C) Scatter plot and linear regression with 95% CI shading of unadjusted psoas major density with age. (D) Bar plot of mean ± SD error bars for adjusted–unadjusted psoas major density.
